# Active geometrodynamics predicts the emergence of cytokinesis

**DOI:** 10.1101/2025.11.07.683232

**Authors:** Aditya Singh Rajput, Kunal Kumar, Masatoshi Nishikawa, K. Vijay Kumar

## Abstract

Cell division accomplishes the segregation of genetic material and involves remarkable changes in the cellular geometry culminating in cytokinesis: the cleavage of a mother cell giving rise to two daughter cells. Cytokinesis in animal cells is driven by flows resulting from cortical tension gradients in the actomyosin cortex. Here, we combine a theory for the active geometrodynamics of the cortical surface and quantitative measurements in the *C. elegans* zygote to reveal the physical principles of cytokinesis. At high activity, we observe the spontaneous emergence of ring-like patterns of myosin concentration and cell shape in the theory. The constriction dynamics of this self-organized pattern agrees with the ingression of the cytokinetic furrow and concomitant myosin accumulation during the first division in the *C. elegans* embryo. Through RNAi perturbations, we quantitatively test our theoretical predictions of myosin accumulation rates linearly varying with the ingression rate, and the emergence of asymmetric ingression. Our work suggests that the self-organised geometrodynamics of active fluid surfaces underlies cytokinesis.

---

The living world is replete with phenomena involving dramatic changes in shape and size at the cellular, developmental, and evolutionary scales (Thompson, 1917; Gerhart and Kirschner, 1997). In animal cells, a prominent morphogenetic process is cytokinesis – the deformation of a mother cell and its subsequent cleavage into two daughter cells (Rappaport, 1996; Forgacs and Newman, 2005). Such shape changes result from flows in the actomyosin cortex located underneath the cell surface (Bray and White, 1988). The shape of the cell affects the nature of the cortical flows and hence the resulting spatial patterns of myosin concentration (White and Borisy, 1983). It is clear that there must exist feedback between the spatial redistribution of cortical tension patterns driven by flows and cellular shape changes. However, the nature of this feedback remains elusive.

The actomyosin cortex is a thin (∼ 0.5 *µ*m) meshwork of cytoskeletal actin filaments, myosin molecular motors, and cross-linking proteins. Myosin motors use the energy of ATP hydrolysis to actuate actin filaments, leading to the generation of contractile stresses in the cortex. This transduction of chemical energy from ATP hydrolysis into mechanical work makes the actomyosin cortex an active material (Jülicher *et al*., 2007; Marchetti *et al*., 2013; Ramaswamy, 2017). Continuous polymerisation and depolymerisation of its constituent elements relaxes elastic stresses in the cortex, with a relaxation timescale ∼ 5 s (Mayer *et al*.; Saha *et al*., 2016). As such, on the timescales of cytokinesis (∼ 10^2^ s), the actomyosin cortex can be regarded as an active viscous thin film. Gradients in these active contractile stresses drive large-scale cortical flows (Mayer *et al*.).

Cortical flows on the cell surface advect myosin motors thereby changing their spatial distribution, which in-turn affects the surface flows via cortical tension gradients. This mechanochemical feedback leads to emergence of self-organized patterns (Bois *et al*., 2011). The same contractile stress patterns generate flows that are orthogonal to the surface of the cell, and hence change its shape (Greenspan, 1977; Salbreux *et al*., 2012). However, the dynamically changing cell shape will affect both the flows and myosin patterns. Can the resulting *geometrodynamic feedback* between the spatial profiles of stress and flow, and cell shape lead to the spontaneous emergence of cytokinesis-like patterns? Can we get experimentally testable predictions for the ingression kinetics from an active geometrodynamic theory for the cell surface? We further ask if the resulting self-organized dynamics can quantitatively account for the accumulation of myosin in the cytokinetic ring and the constriction of the cytokinetic furrow as seen in the cleavage of the *C. elegans* zygote.

In this work, we first set up the geometrodynamics of an active viscous, two-dimensional surface without assuming azimuthal symmetry. The velocity arising from force- and torque-balance conditions advects the concentration of myosin motors and also changes the shape of the surface. Next, we demonstrate that numerical solutions of our model equations lead to the emergence of self-organised ring-like patterns akin to cytokinesis.

Then, we show that the predictions of the theory agree with measurements of the ingression kinetics and the rate of myosin accumulation in the ring. Finally, we show that asymmetries in the ingression of the furrow arise naturally from our theory, and compare well with measurements. In the discussion, we argue that the active deformable nature of the cortex and its ability to spontaneously break azimuthal symmetry quantitatively captures the essential features of cytokinesis.

## GEOMETRODYNAMICS OF THE CELL SURFACE

We model the membrane-cortex composite as a closed and orientable surface embedded in the three-dimensional Cartesian space as shown in FIG. 1(a). At each point on this surface, labelled by the position vector **x**, we have a unique outward unit normal vector 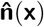. The projection operator 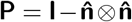, where **I** is the identity tensor and ⊗ denotes the Cartesian tensor product, allows us to identify the tangent plane to the surface at every point. Thus, the velocity vector field **v**(**x**) on the surface can be split into its tangential and normal components as **v** = **v**_∥_ +**v**_⊥_, where 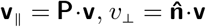, and is the Cartesian dot product (see FIG. 1**a**). Further, denoting the Cartesian gradient by ∇_3_, the surface derivative of a scalar field *c*(**x**) is ∇*c* = **P** · ∇_3_ *c* and the covariant derivative of a vector field is ∇**v** = **P** · (∇_3_ **v**) · **P** (Delfour and Zolésio, 2011; Walker, 2015). The shape operator 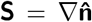 encodes the extrinsic curvature of the surface which describes the spatial variation of 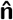 (see FIG. 1**a**). From **S**, we obtain the summed curvature ℋ = Tr(**S**) and the Gaussian curvature 𝒦= [ℋ^2^ − Tr(**S**^2^)]*/*2, where Tr denotes the trace operation.

**FIG. 1.**
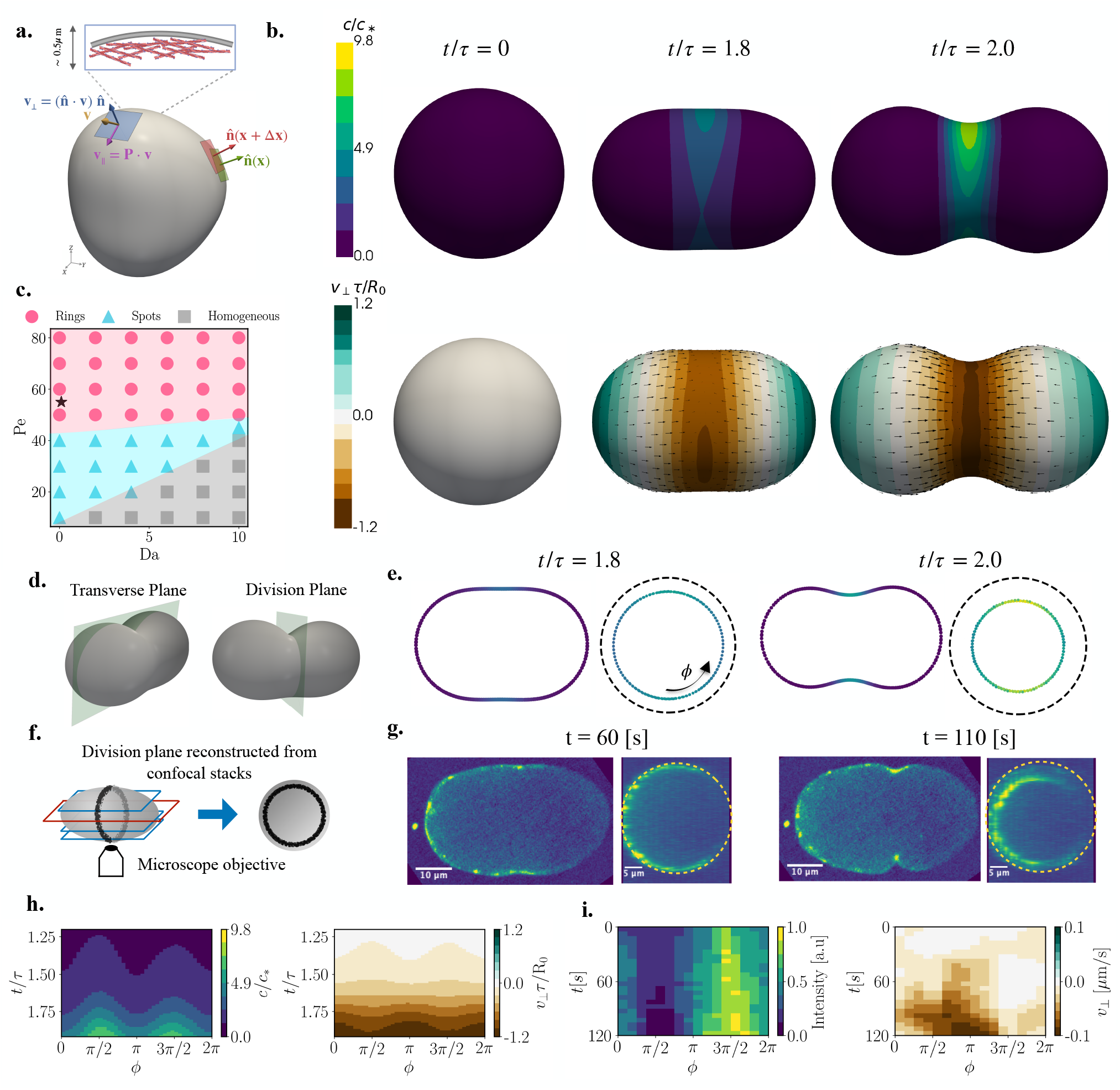
Emergence of ring-like patterns. **a**. The actomyosin cortex (thickness ∼ 0.5*µ*m), along with the cell membrane, is considered as a two-dimensional active fluid surface. Gradients of surface active stresses lead to the hydrodynamic velocity field. **v**. The normal component 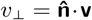 deforms the surface while the tangential component. **v**_∥_ =. **P·. v** advects surface concentration fields. Here 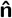 is the unit-outward normal to the surface and 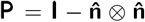 is the tangential projector. **b**. Time sequence of the surface shape, along with myosin concentration in the top row, surface deformation rate and the tangential velocity. **v**_∥_ in the bottom row, show the spontaneous emergence of a cytokinetic furrow like pattern. Time here is measured in units of *τ* = *η/*(*γD*). **c**. Phase-diagram of emergent patterns in the contractility (Pe) and the turnover-rate of myosin (Da) show that spherical shapes with a homogeneous myosin distribution are seen at low Pe. Upon increasing Pe, we see the formation of spot-like patterns. And at even higher Pe furrow-like patterns of surface shape and ring-like patterns of myosin are seen. Here Λ = 1*/*2 and ℬ= 1*/*50. The time sequence shown in. **a** corresponds to the ⋆ marked in. **c. d**. For further analysis, we view these patterns in the division plane and the transverse plane. **e**. Cross-sections of the surface in these two planes show the development of the furrow. As in. **b**., the colour indicates the myosin concentration field. **f**. Schematic of fluorescence image acquisition of NMY-2::GFP and reconstruction of the division plane from the confocal stacks in *C. elegans* zygotes. **g**. The cell shape and myosin patterns seen in the transverse and division planes are qualitatively similar to the theoretical predictions in. **e**. Kymographs of the myosin concentration (left) and the ingression rate (right) in the division plane from the theory (**h**) compare well with those extracted from experiments (**i**).

### Kinematics of fluid-surfaces

The actomyosin cortex flows like a fluid on timescales longer than the relaxation time for the elastic stresses (Saha *et al*., 2016). Given a surface velocity field **v**(**x**) on such a “fluid-surface”, the movement of a material point **x** is governed by the normal component of **v**, i.e.,

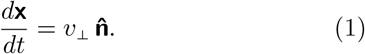

As the surface deforms, its geometry changes. The surface gradient of *v*_⊥_ tilts the local tangent plane. In other words, the evolution of 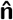 satisfies (Waxman, 1984; Salbreux and Jülicher, 2017; Jankuhn *et al*., 2018):

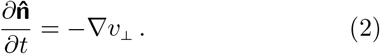

The evolution equations for **P** and **S** follow from the above equation (see SI). Equations (1) and (2) govern the kinematics of a fluid surface for a given velocity **v**, which must be obtained from the mechanics of the surface.

### Surface velocity from force-balance

The internal forces arising from interactions between the fluid-elements on the surface are captured by a stress tensor **Σ**. On cellular scales, inertial forces are negligible. As such, the velocity field **v** is obtained from force- and torque-balance conditions encapsulated by ∇ · **Σ** = −**F**_ext_, where **F**_ext_ is the total external force on the surface. We neglect hydrodynamic flows in the cytoplasm, approximate its dissipative effects by a frictional force −*γ* **v** (Salbreux *et al*., 2009), and incorporate its incompressibility by including a constraint force 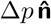, where the Lagrange multiplier Δ*p* is the spatially uniform pressure difference across the surface obtained by implementing the volume constraint ∫_surface_ *v*_⊥_ = 0. The total external force is thus 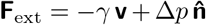. To hold the centre-of-mass fixed, we include an additional force 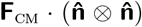, where **F**_cm_ is a time-dependent Lagrange multiplier implementing the constraint 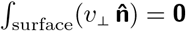.

The hydrodynamic stress **Σ** has contributions from the viscous stresses arising from in-plane velocity gradients ∇**v**, the forces arising from the bending elasticity of the surface, and finally the nonequilibrium active stresses arising from the ATP consuming activity of molecular motors. Assuming the surface behaves as a Newtonian fluid, the viscous stress is **Σ**_viscous_ = 2*η* **E** + (*η*_*b*_ − *η*) Tr(**E**) **P**, where *η* and *η*_*b*_ are, respectively, the shear and bulk viscosities, and the surface strain-rate tensor **E** = [∇**v** + (∇**v**)^T^]*/*2 with (· · ·)^T^ denoting the transpose. Note that we have assumed that cytoplasmic exchange maintains the material density of the surface at a constant value, and have also neglected the two-dimensional pressure (Jülicher *et al*., 2018). The Canham-Helfrich energy *ℰ*_ch_ = *β ∫*_surface_ *ℋ*^2^, where *β* is an elastic modulus, governs the bending elasticity of a symmetric bilayer without any spontaneous curvature (Canham, 1970; Helfrich, 1973). The equilibrium shapes of such fluid-surfaces, corresponding to the minima of *E*_CH_, can be obtained by considering the bending stress 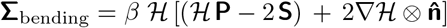(Salbreux and Jülicher, 2017; Capovilla and Guven, 2002; Capovilla *et al*., 2005) (see SI). Thus, the geometrodynamics of a passive fluid-surface is governed by Equations (1) and (2), where the velocity is obtained by solving ∇ *·* (**Σ**_viscous_ + **Σ**_bending_) = −**F**_ext_.

### Active deformable surfaces

The active contractile stress **Σ**_active_ in the cortex is locally regulated by the concentration *c*(**x**) of myosin motors. Assuming that this active stress is isotropic in the tangent plane of the surface, we consider **Σ**_active_ = *ζ* Δ*µ f* (*c*(**x**)) **P**, where *ζ >* 0 is a proportionality constant, Δ*µ >* 0 is the difference in chemical-potential associated with the hydrolysis of ATP, and the active-stress regulation function *f* (*c*) increases monotonically with *c* (Mayer *et al*.; Bois *et al*., 2011; Kumar *et al*., 2014; Gross *et al*., 2019; Mietke *et al*., 2019a,b). Without loss of generality, we take *f* (*c*) = *c/*(*c* + *c*_∗_), with *c*_∗_ being a saturation concentration. Therefore, the velocity field **v**(**x**) on the active fluid-surface is obtained by solving

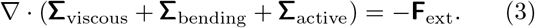

This velocity field has two consequences: first, its normal component changes the shape of the surface via (1) and, second, it advects the concentration of myosin motors on this deforming surface. The concentration field *c*(**x**) then evolves according to

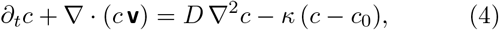

where *D* is an effective diffusion constant, *κ* is a turnover rate, and *c*_0_ is the set-point concentration of the turnover reaction. To conclude, the geometrodynamics of the active fluid surface is specified by the Equations (1)-(4).

Given an initial shape (say a sphere of radius *R*_0_) and an initial concentration field on the surface, the Equations (1)-(4) completely determine the subsequent spatiotemporal evolution of both the surface geometry and the concentration patterns. This dynamics is characterised by the following dimensionless parameters: the Péclet number Pe ≡ *ζ*Δ*µ/*(*γD*) compares active- and diffusive-transport, the Damköhler number 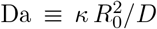 compares the turnover rate with a diffusion timescale, a non-dimensional hydrodynamic length-scale 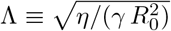, and a non-dimensional bending rigidity *ℬ* ≡*β/*(*γ D*). In our theoretical results, we report concentration and time in units of *c*_∗_ and *τ* = *η/*(*γ D*). For this study, we fix *c*_0_*/c*_∗_ = 1 and *η*_*b*_*/η* = 5*/*4, and start with a spherical surface of radius *R*_0_, and a uniform concentration of *c* with small random fluctuations. Note that even though the velocity **v** is determined by the physical properties of the surface, Equations (1) and (2) determining the kinematics of the surface geometry do not have any material parameters.

## RESULTS

We ask what kinds of surface geometries and concentration patterns result from the self-organized dynamics of our model Equations (1)-(4). To solve these equations in the non-linear regime without assuming azimuthal symmetry of either the surface geometry or the concentration patterns, we developed a robust and flexible numerical simulation based on the finite-element-method using the FEniCS project (Alnæs *et al*., 2015) (see SI). Clearly, a spherical shape and a uniform concentration *c*(**x**) = *c*_0_ (with a concomitant velocity **v**(**x**) = **0**) is a possible solution. Using spherical harmonics 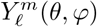 where 𝓁 = 0, 1, …, −𝓁 ≤ *m* ≤ 𝓁, *θ* and *φ* are respectively the polar and azimuthal angles, a linear stability analysis reveals that this state can become unstable at large Pe (Mietke *et al*., 2019a). The fastest growing mode beyond the stability-boundary of the homogeneous state is the 𝓁 = 1 mode, and our numerical simulation recovers this stationary patterned state (see SI).

### Spontaneous emergence of cytokinetic ring-like patterns

At larger values of Pe, we see the spontaneous emergence of cytokinesis-like patterns in both the surface geometry and the myosin concentration field *c*(**x**). As shown in the top row of FIG. 1**b**, an initially uniform concentration field with small random perturbations starting from spherical shape spontaneously develops a ring-like pattern of myosin in time. Concomitantly, the geometry of the surface deforms to a shape resembling a cytokinetic furrow. The bottom row of FIG. 1**b** shows the spatiotemporal evolution of the active flow **v**(**x**) on the deforming surface: the glyphs represent the tangential velocity **v**_∥_ while the heat-map represent the normal velocity *v*_⊥_. It is well known that at large Pe, inhomogeneities in myosin concentration can accumulate to destablise the homogeneous state on fixed domains (Bois *et al*., 2011). However, in our case, the surface also deforms in response to the velocity field associated with these concentration patterns.

In FIG. 1**c**, we show a phase-diagram of the patterns obtained from our theoretical model as we vary the strength of the active stresses (Pe) and the turnover-rate (Da). At low Pe, the state with a homogeneous concentration and a spherical shape is the stable solution. Upon increasing Pe, we find a spot-like pattern of the concentration and a concomitant deformed shape as the stable solution. At larger Pe, this state undergoes an instability wherein the concentration field self-organizes in the form of a ring-like pattern and the shape ingresses continuously. Note that this latter state is not a steady-state solution. Eventually, the surface ingresses to a large extent leading to self-intersections in our numerical implementation, at which point our theory is no longer valid. However, we can follow the kinetics of this ingression process starting from a spherical shape. We observe that during this process, the ring remains planar and is approximately circular. Varying the hydrodynamic length Λ and the bending rigidity ℬ, we find that a broad range of parameters can lead to the emergence of ring-like patterns of concentration and the concomitant furrow-like shapes of the surface (see SI).

In FIG. 1**d**, we define the transverse and the division planes, and in FIG. 1**e** we show the surface shape at two timepoints along with the concentration values (indicated by a colormap) in the transverse and division planes. We immediately notice that the surface ingresses asymmetrically: we will return to this point later. To conclude, our theoretical model shows the spontaneous emergence of a cytokinetic ring-like patterns in both the concentration field and the geometry of the surface.

### Cytokinesis in C. elegans zygotes

To compare the predictions of our theoretical model with experimental data, we consider the first embryonic division in *C. elegans*. We imaged GFP-labelled non-muscle myosin NMY-2 using confocal microscopy (FIG. 1**f**). Combining several Z-stacks, we obtain the three-dimensional spatiotemporal dynamics of NMY-2::GFP (see SI movie). In FIG. 1**g**, we plot the cell shape and myosin intensity in the transverse and division planes, at two different time points, as seen in *C. elegans* zygotes. Notice the remarkable similarity in FIG. 1**e** and FIG. 1**g** of both the concentration and shape patterns. In FIG. 1**h**, we show the kymograph of the concentration pattern and the ingression rate (*v*_⊥_) in the division plane from our numerical calculations. We compare this with the kymographs of the NMY-2::GFP and normal velocity of the contractile ring seen in FIG. 1**i**.

### Kinetics of ingression

Next, we follow the kinetics of the ingression process once a ring-like pattern emerges. To do so, we look at the following quantities defined in the division plane (see FIG. 2**a**): (i) the Gaussian curvature *K* = *k*_1_ *k*_2_ where *k*_1_ and *k*_2_ are the principal curvatures of the surface, (ii) the furrow radius *R*(*t*), and (iii) the concentration of the stress-regulator in the ring. First, in FIG. 2**b**, we plot the average Gaussian curvature in the ring 𝒦^ave^(*t*) = (∫_ring_ 𝒦) */*(2*πR*(*t*)) from our theory. We notice that 𝒦^ave^ decreases from an initial positive value to negative values (indicating the development of a furrow-like shape) at later times. Further, we see that the rate at which 𝒦^ave^ decreases is an increasing function of the activity, i.e., Pe. To obtain the corresponding data for the *C. elegans* embryo, we performed fluorescence imaging of a GFP-fused PH domain bound to the membrane (see SI). This allows us to digitally reconstruct the cell surface as a triangulated mesh, and extract the Gaussian curvature (Cohen-Steiner and Morvan, 2003). In FIG. 2**c**, we see the temporal evolution of the average Gaussian curvature 𝒦^ave^ in the wild-type (WT) embryo. We notice that 𝒦^ave^ is almost zero at initial times, and decreases towards large negative values at later times indicating the development of the furrow. Note that the negative values of 𝒦^ave^ at the onset of ingression are due to low signal-to-noise ratio. To test the theoretical prediction about the rate of decrease of 𝒦^ave^, we performed RNAi to modulate the contractility of the cortex, which affects the strength of the cortical flows (Naganathan *et al*., 2018). We see in FIG. 2**c** that *gsp-1* RNAi embryos (which display enhanced cortical flows) show a faster rate of decrease of 𝒦^ave^ while this rate decreases in *nmy-2* RNAi embryos (which have reduced cortical flows) compared to WT.

**FIG. 2.**
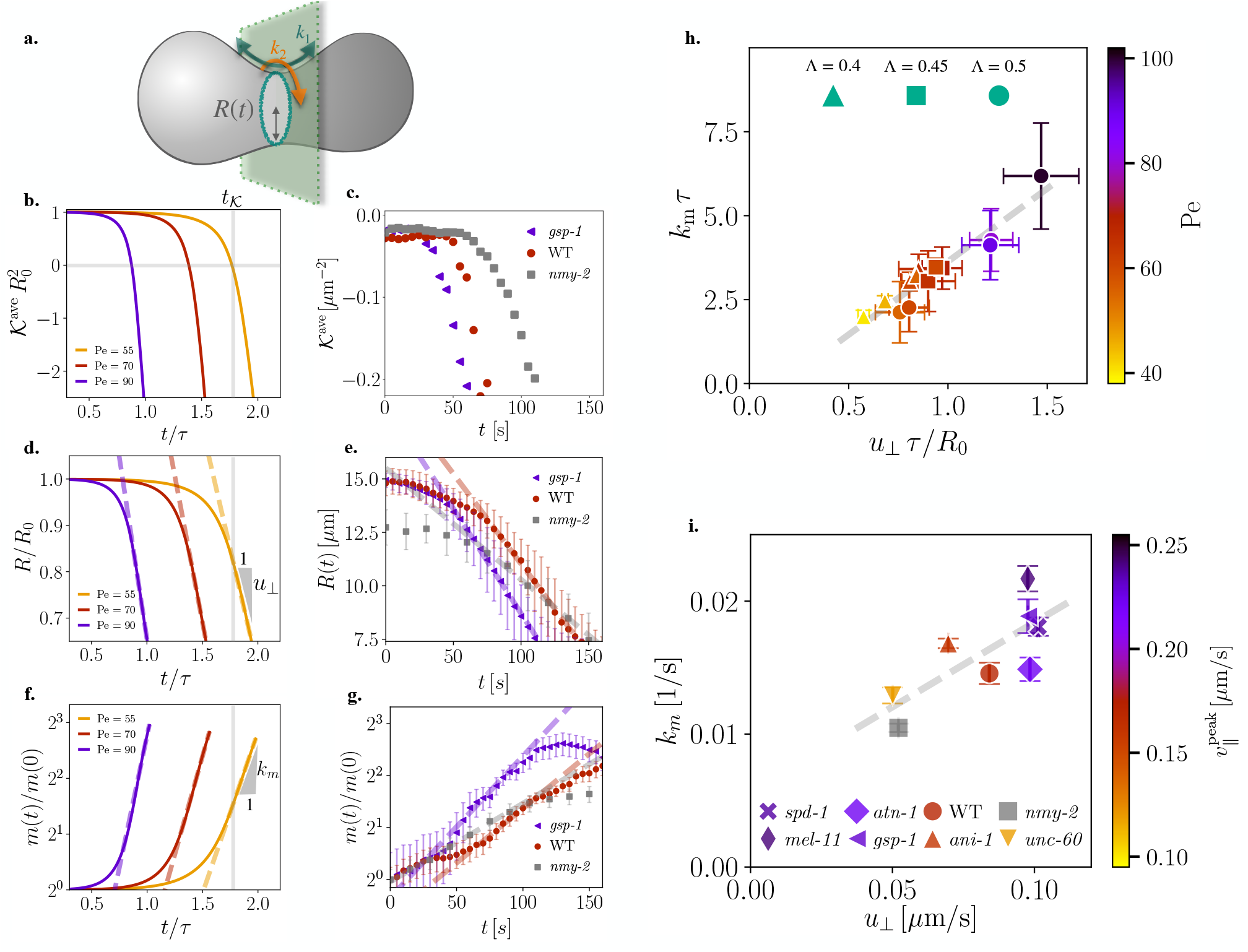
Geometrodynamics of cytokinesis. **a**. In the division plane, *R*(*t*) denotes the radius of curvature of the furrow. The Gaussian curvature of the surface is 𝒦 = *k*_1_ *k*_2_, where *k*_1_ and *k*_2_ = 1*/R* are the principal curvatures of the surface. **b**. The ring-averaged Gaussian curvature 𝒦^ave^ starts from positive values and crosses zero at *t* = *t*_𝒦_ when the surface starts developing a furrow-like region. The rate of decrease of 𝒦^ave^ increases with Pe. **c**. Reconstructing the cell surface using a membrane marker (PH::GFP), we extract the Gaussian curvature. Compared to the wild-type (WT) *C. elegans* embryo, the strength of cortical flows is higher in *gsp-1* RNAi embryos and lower in *nmy-2* RNAi embryos. We observe that 𝒦^ave^ decreases in a concomitant manner. **d**. The radius *R*(*t*) of the furrow decreases linearly around *t* = *t*𝒦 and the rate of decrease *u*_⊥_ = − *dR/dt* increases with Pe. **e**. These features are also observed in *C. elegans* embryos. **f**. The ring-averaged myosin concentration 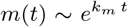 shows an exponential increase around *t* = *t*_𝒦_. **g**. We observe this exponential increase across RNAi conditions. **h**. We plot *k*_*m*_ versus *u*_⊥_ over a range of Pe and Λ, and observe a linear relation between them as predicted by our theory in (5) and (6). Here the errorbars are standard error over random initial conditions. **i**. We extract *u*_⊥_ and *k*_*m*_ from the fits (dashed-lines) in. **e** and. **g** respectively and observe that *k*_*m*_ is linear in *u*_⊥_. We use the peak cortical speed 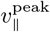 as a measure of the contractility. In. **b, d**, and. **f**, Da = 1*/*10, Λ = 1*/*2 and ℬ= 1*/*50, while in. **h**, Da = 1*/*10 and ℬ= 1*/*50. The error bars in. **e** and. **g** are standard error of the mean with *N* = 11 (WT), *N* = 5 (*spd-1*), *N* = 7 (*mel-11*), *N* = 5 (*atn-1*), *N* = 11 (*gsp-1*), *N* = 6 (*ani-1*), *N* = 11 (*nmy-2*), *N* = 7 (*unc-60*).

Second, FIG. 2**d** shows the evolution of the furrow radius *R*(*t*) as a function of time from our theory. We notice that after an initial slow phase, *R*(*t*) decreases linearly with time, thus defining the constant ingression rate *u*_⊥_ ≡ −*dR*(*t*)*/dt*. This rate is related to the ring-averaged normal velocity of the surface ∫_ring_ *v*_⊥_(*t*). FIG. 2**e** compares the evolution of *R*(*t*) for three different RNAi conditions in the *C. elegans* embryo and shows that *R*(*t*) decreases linearly with time, consistent with the theory. This linear decrease in the furrow radius as a function of time seems to be a universally observed feature across several eukaryotes (Mabuchi, 1994; Pelham Jr and Chang, 2002; Biron *et al*., 2005; Zumdieck *et al*., 2007; Carvalho *et al*., 2009; Khaliullin *et al*., 2018).

Third, we follow the dynamics of the ring-averaged concentration *m*(*t*) = (∫ _ring_ *c*(*t*)) */*(2*πR*(*t*)) in FIG. 2**f**, and notice that it increases exponentially with time 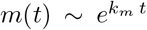, thus defining the rate *k*_*m*_. In FIG. 2**g**, we plot the NMY-2::GFP intensity across different RNAi conditions in the *C. elegans* embryo and notice that it shows an exponential increase as predicted by our theory. This observation is in line with previous studies (Khaliullin *et al*., 2018).

To understand the linear decrease in the furrow radius and the exponential increase of myosin in the ring, we derive simplified equations for the dynamics of *R*(*t*) and *c*(*t*) using some reasonable approximations around the time-point *t*_𝒦_ (see FIG. 2**b**) where the average Gaussian curvature changes sign (see Methods). For small *t* − *t*_𝒦_, we find

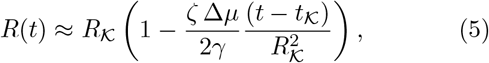

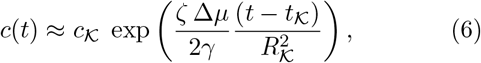

where *R*_𝒦_ and *c*_𝒦_ are the values of the radius and the (uniform) concentration at *t* = *t*_𝒦_. The above expressions show that active contractility underlies both the dynamics of shape change and the accumulation of myosin in the cytokinetic furrow. In fact, we predict a linear relationship *k*_*m*_ ∼ *u*_⊥_*/R*_𝒦_. However, this might not hold exactly due to our approximations.

As expected, we notice from FIG. 2**b** and **f** that the rates *u*_⊥_ and *k*_*m*_ increase with Pe. We ask if there is a linear relationship between them as predicted above. To answer this, we numerically solved our model Equations (1)-(4) for a range of Pe values, at different values of the turnover rate Da, and the hydrodynamic length-scale Λ, and averaged over random initial conditions. In FIG. 2**h**, we plot *k*_*m*_ against *u*_⊥_, where the colour of the symbol maps to different values of Pe, and observe a linear relation as predicted by the above approximate calculation.

We next ask if *k*_*m*_ and *u*_⊥_ are correlated during cytokinesis in the *C. elegans* zygote. We notice from FIG. 2**c** and **g** that the rates *u*_⊥_ and *k*_*m*_ vary across RNAi conditions in the embryo. To quantify the relationship between these rates, we first measured the in-plane cortical flows in the embryo and use their peak value 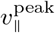 as a direct measure of activity (see SI). In FIG. 2**i**, we plot *k*_*m*_ against *u*_⊥_ extracted from the data in the embryo for different RNAi conditions, where the colour of the symbol now maps to the strength of 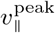. As predicted by our theory, we see a positive linear correlation between *k*_*m*_ and *u*_⊥_. To conclude, the predictions of our theoretical model incorporating the active geometrodynamics of the surface for the self-organized kinetics of ingression agrees with experimental measurements. In other words, these results imply that the shape changes during cytokinesis, while initiated by signals, is possibly a self-organized process driven by actomyosin contractility.

### Emergent asymmetries during cytokinetic ingression

We next turn to the asymmetric ingression of the surface that we alluded to earlier (Maddox *et al*., 2007; Hsu *et al*., 2023). Both the concentration pattern of the active stress regulator and the surface geometry seen in FIG. 1**b**, as well as the spatial maps of NMY-2::GFP and the cross-section of the cell seen in FIG. 1**g**, show that there are asymmetries during ingression. This is also clearly seen in the kymographs in FIG. 1(**h-i**). As shown in FIG. 3**a**, the deviation Δ of the center of the cytokinetic ring from its initial position, is a quantitative measure of this asymmetry (Menon *et al*., 2017). In FIG. 3**b**, we plot the temporal evolution of Δ and notice that it increases exponentially, i.e., 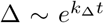 which defines the rate-constant *k*_Δ_. FIG. 3**c** shows the variation of Δ with time extracted from the embryo for several RNAi conditions. As predicted by our theory, Δ increases exponentially with time. We further note that *k*_Δ_ increases with Pe in the theory and with the strength of the cortical flows in the cell. We now ask if there is a correlation between the rate *k*_Δ_ at which the asymmetry increases and the ingression rate *u*_⊥_ of the furrow. In FIG. 3**d**, we plot *k*_Δ_ versus *u*_⊥_ from our theory, where the colour of the symbols map to the value of Pe. Correspondingly, we plot in FIG. 3**e** *k*_Δ_ versus *u*_⊥_ extracted from the embryo across different RNAi conditions, where the colour of the symbols now represent 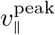 as defined earlier. We clearly see both from theory and experiment that the rate at which the cytokinetic furrow ingresses asymmetrically is linearly correlated with the ingression rate of the furrow. To conclude, our theoretical model for the self-organized geometrodynamics of the surface not only predicts the ingression of the cytokinetic furrow, but also predicts the concomitant asymmetric ingression, in good comparison with experimental data.

**FIG. 3.**
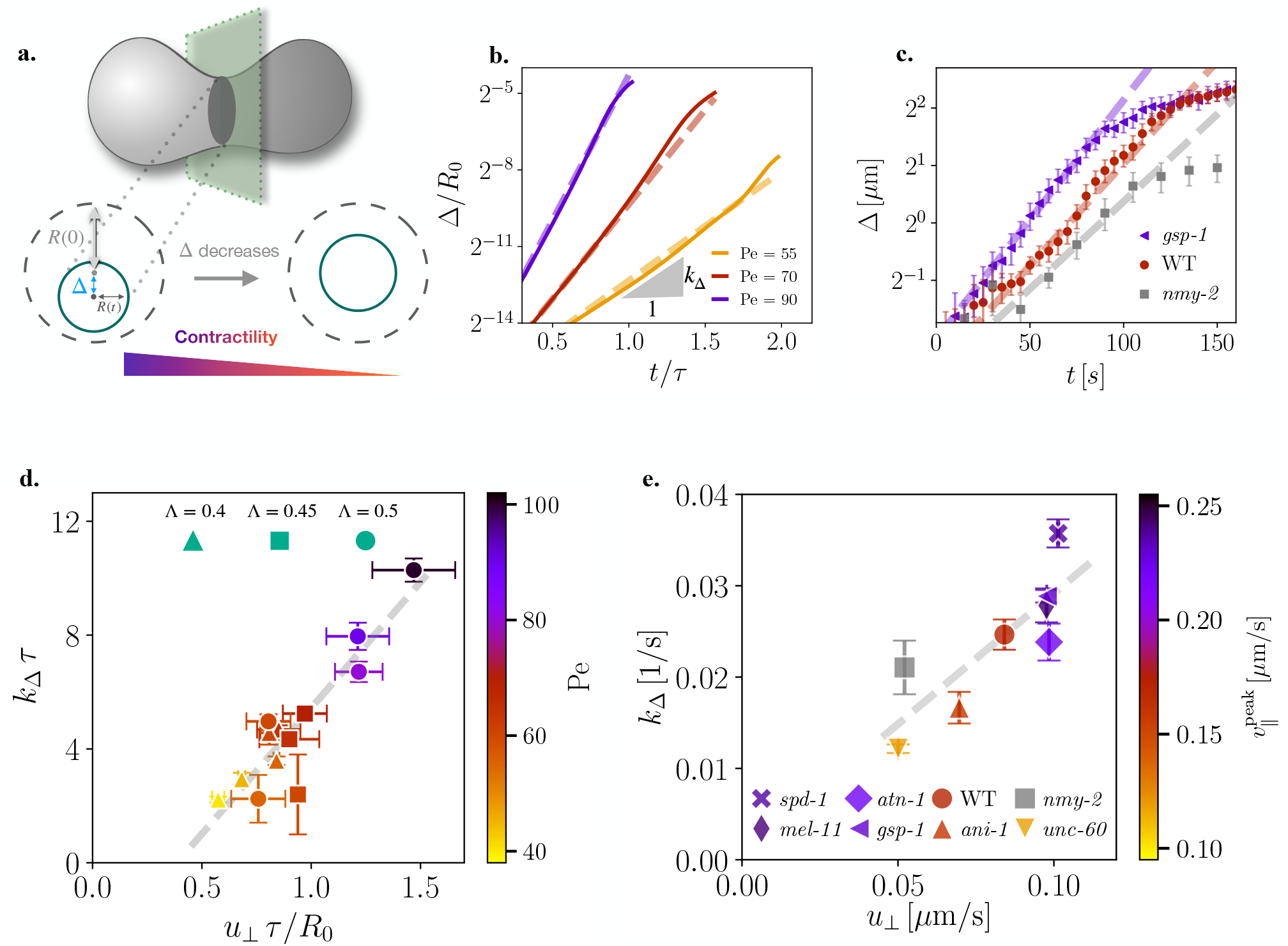
Asymmetries in cytokinetic ingression. **a**. As cytokinesis proceeds, the center of the cytokinetic ring does not remain fixed. The displacement Δ of the center of the ring from its location at *t* = 0 is a measure of the asymmetry of ingression. The asymmetry Δ reduces upon decreasing contractility. **b**. From our numerical simulations, we predict an exponential increase 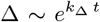 of the asymmetry with the rate *k*_Δ_. Furthermore, we predict *k*_Δ_ increases with Pe. **c**. During cytokinesis in the *C. elegans* zygote, we observe an exponential increase of Δ. Compared to WT embryos, measured rate *k*_Δ_ is higher for *gsp* − 1 RNAi embryos while it is lower for *nmy* − 2 RNAi embryos. **d**. In our numerical simulations, we find *k*_Δ_ increase linearly with the ingression rate *u*_⊥_. **e**. We find that this feature is also observed during the first division of the *C. elegans* embryo across RNAi conditions. Taken together,. **d** and. **e** imply that embryos with higher contractility (higher 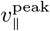) show higher asymmetry during ingression. The parameter values and the calculation of errorbars are as in FIG. 2.

### Summary of results

We summarize our findings by noting the following points.

First, in our theoretical model, we hypothesized that the active stresses responsible for tangential flows on the surface are also the sources for the normal flows that deform the surface. The dynamical geometry of the surface, in turn, has a strong feedback on the patterns of the active stress regulators. We have shown that this tight coupling between active patterns and dynamical shapes – active geometrodynamics – leads to spontaneous formation of cytokinesis like patterns.

Second, the agreement between the theoretical model and experimental data from the *C. elegans* embryo for the ingression kinetics strongly suggests that the underlying physics of cytokinesis is governed by the large-scale self-organized nature of the chemical and shape patterns. In particular, the predictions of the linear decrease in the furrow radius, the exponential increase in the concentration of the active-stress regulator in the ring, and the exponential increase in the asymmetry of the ring that emerge naturally in our theoretical model are confirmed by the data. Moreover, the relationship between the kinetic rates (*u*_⊥_, *k*_*m*_, and *k*_Δ_) predicted by the theory across various parameter values are borne out in the experiment across different RNAi conditions.

Third, consistent with the predictions of a linear stability analysis of the homogeneous state, neglecting cytoplasmic flows (Mietke *et al*., 2019a; Picardo *et al*., 2025), the initial patterns of concentration are localized spots, even in the regime where we find rings. However, the non-linearities in the active geometrodynamics of the surface lead to the merger of these spot-like patterns, resulting in the emergence of a ring-like pattern.

Fourth, though these localized spots merge to form a ring-like pattern, the concentration is not uniform in the ring. Moreover, the positive geometrodynamic feedback between concentration patterns and the surface shape amplifies the inhomogeneities of the concentration in the ring, leading to asymmetries in ingression. In our theory, we find that the enhancement of these inhomogeneities can occur when the active flows are sufficiently strong. In other words, the timescale for the growth of inhomogeneous patterns resulting from active flows should be smaller than the timescales for the homogenization of the concentration by isotropic process such as diffusion and turnover (Thiyagarajan *et al*., 2022).

Fifth, in the cell, the initiation of cytokinesis is under tight control of the cell-division cycle. The signals for this initiation can be localized in space and time (for instance, through positioning of the spindle) or lead to global changes in material properties (such as contractility) of the cortex at appropriate temporal checkpoints (Fischer-Friedrich *et al*., 2016). However, our study suggests that once cytokinesis is initiated, the subsequent spatiotemporal patterns of the active-stress regulators and the geometry of the cell surface could result from a self-organized dynamics, without the continued presence of signals. For instance, it is known that chemical or mechanical perturbations to the spindle at appropriate times do not hamper the cell’s ability to undergo cytokinesis (Swann and Mitchison, 1953; Hiramoto, 1956).

## DISCUSSION

The minimal theoretical model for the geometrodynamics of the surface considered in this study neglects several realistic features of the cell. First, comparing Fig. 1 **e** and **g**, we notice that the cytokinetic furrow has a high curvature in the *C. elegans* embryo compared to the theoretically predicted shape. This contrast possibly arises because the embryo is confined in an eggshell, which prevents the daughter cells from moving apart. A few studies indicate that removing this eggshell leads to a lower curvature in the ingression furrow (Koyama *et al*., 2012).

Second, in our theory, we have neglected the nematic alignment of actin filaments in the cytokinetic ring (Reymann *et al*., 2016; Spira *et al*., 2017) and have considered an isotropic active stress in the tangent plane of the surface. Our experiments in the embryo show that the tangential component of the cortical flows towards the ring decreases after the onset of cytokinesis (see SI). However, our theory predicts a monotonic increase in the magnitude of the tangential flows **v**_∥_, well beyond the onset of furrow ingression. It is possible that including the dynamics of the orientational degrees of freedom of the actomyosin cortex, and the associated anisotropic active stresses, might reduce the tangential flows from the poles towards the ring.

Third, previous theoretical studies have shown that including cytoplasmic flows can lead to the emergence of ring-like patterns of the concentration field (Greenspan, 1977; Mietke *et al*., 2019b). However, this mechanism requires a strong coupling between the cortical and cytoplasmic flows, with ring-like patterns emerging when *η/*(*µ R*_0_) ≪1, where *µ* is the cytoplasmic shear viscosity. Using values of the cortical shear viscosity obtained in other cells (Turlier *et al*., 2014) and considering appropriate values for the shear viscosity of the cytoplasmic fluid (Charras *et al*., 2009), we estimate *η/*(*µ R*_0_) ∼ 10. This indicates that ring-like patterns are unlikely to emerge from a strong coupling between cortical and cytoplasmic flows.

Fourth, this study has focused on the asymmetry of the cytokinetic ingression furrow and has not focused on asymmetric cell division wherein the two daughter cells are different in size and/or chemical-composition. Asymmetric cell division is typically preceded by the establishment of cell-polarity (Gross *et al*., 2019). It is not clear if there exist correlations between asymmetric cell division and asymmetric ingression. This aspect needs to be investigated further by explicitly including cell polarity.

Fifth, the self-organised nature of cytokinetic ingression that we have demonstrated indicates that cytokinesis-like patterns could arise in synthetic systems (Baldauf *et al*., 2022). Moreover, since our work demonstrates that the physics of active deformable surfaces underlies cellular morphogenesis, we expect similar principles to underlie shape dynamics at larger scales, such as tissues, wherein gradients of myosin activity can drive tissue morphogenesis (Streichan *et al*., 2018; Gehrels *et al*., 2023).

Our study indicates that geometrodynamics of an active surface can lead to the spontaneous emergence of cytokinesis-like patterns. The theoretical model naturally predicts the emergence of spontaneous asymmetries during ingression. We find that these self-organised patterns of concentration and shape are in agreement with those seen in the *C. elegans* embryo. Moreover, the kinetics of concentration accumulation in the furrow and the surface dynamics, including asymmetric ingression, agree with experimental measurements across different RNAi conditions and previous studies. It is remarkable to note that the linear decrease in the size of the cytokinetic furrow predicted by our theory has been observed in many eukaryotic species (Mabuchi, 1994; Pelham Jr and Chang, 2002; Zumdieck *et al*., 2007; Carvalho *et al*., 2009). Furthermore, from both theory and experiments, we have discovered universal relationships among *u*_⊥_, *k*_m_ and *k*_Δ_ (FIG. 2 **h-i** and FIG. 3 **d-e**) during cytokinesis. Taken together, these results reveal the crucial role of dynamical geometry, and its coupling to mechanochemical processes, in the physical principles underlying cytokinesis.

## Acknowledgements

ASR, KK, and KVK acknowledge support of the Department of Atomic Energy, Government of India, under project number RTI4001. MN acknowledges support from Grant-in-Aid for Transformative Research Areas (KAKENHI), 22H05110. KVK acknowledges support from the Max Planck partner group at ICTS-TIFR, the Department of Biotechnology (India) through a Ramalingaswami reentry fellowship, and the financial support of the John Templeton Foundation (grant 62220). This research was also supported in part by ICTS-TIFR for participating in the programs ICTS/amab2023/11 and ICTS/SOBF2025/10. ASR and KVK thank Vidyanand Nanjundiah, Sriram Ramaswamy, Vishal Vasan, Maithreyi Narasimha, and Vikas Trivedi for discussions.

## METHODS

### Experimental methods

#### Worm strains and culture

*C. elegans* strains were maintained at 20°C and shifted to 25°C for 24 hr under nonRNAi condition. RNAi was performed via feeding, as described in (Naganathan *et al*., 2018). Feeding durations were 9-10 hr for *unc-60*, 15-17 hr for *cdc-42*, 16-18 hr for *nmy-2*, and 27-29 hr for *ani-1*. For *atn-1, gsp-1, mel-11*, and *spd-1*, feeding was conducted for 29-31 hr at 25°C. The following transgenic strains were used: LP162 for NMY-2::GFP imaging, and LP306 for imaging GFP-tagged PH domain for plasma membrane imaging.

#### Time-lapse imaging

Z-stack confocal images were acquired using a Nikon Ti2-E microscope with a Yokogawa CSU-W1 spinning disk unit, a NIDAQ-triggered Piezo Z-stage (MCL Nano-Z-100-N2), and a Nikon PlanApo 60× water-immersion objective (N.A. 1.2). Images were captured by an sCMOS camera (Hamamatsu ORCA-Fusion) at 256 *×* 256 pixels, using 2 *×* 2 binning from an original 512 *×* 512 pixel region. GFP-tagged proteins in zygotes were excited with a 488 nm diode laser. For NMY-2::GFP imaging, the laser power was 15 mW (30 mW for *nmy-2* RNAi), with a 40 ms exposure. Z-stacks of 20 slices at 1.5 *µ*m intervals were acquired every 5 s (15 s for *nmy-2* RNAi). For GFP-PH domain imaging, a 4 mW laser with a 20 ms exposure was used; 10-slice Z-stacks at 1 *µ*m spacing were acquired every 5 s.

One-cell embryos were dissected from young adults in M9 buffer and mounted between a coverslip and slide using 25 *µ*m spacer beads (for NMY-2 imaging) or 10 *µ*m beads (for PH domain imaging), sealed with VALAP. Smaller beads were used to bias ingression initiation from the lateral side; larger beads were used to suppress embryo rotation prior to cytokinesis onset.

### Approximate theory for ring kinetics

In this section, we derive an approximate theory for the kinetics of the ring radius and the (average) concentration of myosin in the ring. Around the time-point *t* = *t*_𝒦_ where the (average) Gaussian curvature vanishes and the surface starts developing a neck-like region, we derive approximate equations for the ring radius *R* and the (averaged) myosin concentration *c*. Note that at *t* = *t*_𝒦_, the summed curvature *ℋ* = 1*/R*.

We assume that (i) both the surface shape and the concentration of myosin are azimuthally symmetric in the ring, (ii) the surface variation of the myosin concentration peaks at the location of the ring, (iii) the tangential fluxes of myosin (from advection and diffusion) into the ring balance each other, and (iv) the shape deformations are volume preserving so that Δ*p* ≈ 0. Assumption (i) implies that the myosin concentration is uniform along the ring, assumption (iii) implies **v**_∥_ *c* − ∇*D*∇*c* = **0**, i.e., the concentration in the ring changes only due to the evolution of geometry, while assumptions (ii) and (iv) imply that **v**_∥_ = **0** on the ring. Since we observe ring-like patterns even when *κ* = 0, we neglect myosin turnover.

#### Dynamics of the ring radius

The force- and torque-balance condition in the normal direction takes the form

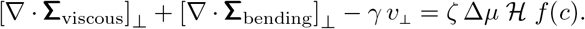

We neglect the forces arising from viscous stresses, and for small deviations of the surface around *t* = *t*_𝒦_, we retain terms to lowest order in curvatures while neglecting curvature gradients in the bending stresses. We then get

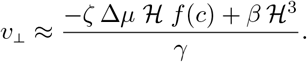

The equation governing the shape evolution leads to

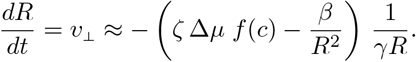

#### Dynamics of the myosin concentration in the ring

The concentration field evolves only due to the changing geometry:

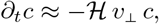

which using the above expression for *v*_⊥_ leads to

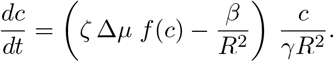

Taken together, the above equations for *R* and *c* is our approximate theory for the ingression kinetics. We emphasize that these equations are valid only for short times around *t* = *t*_𝒦_.

#### Rates of ingression and myosin accumulation

If *c* ≫ *c*_∗_ (which is the case seen in FIG. 1**b**), then *f* (*c*) ≈ 1. In this case,

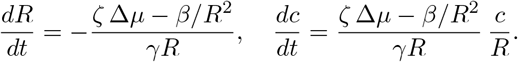

We clearly see that for a weakly active surface, i.e., *ζ* Δ*µ/β* ≪ *R*^−2^, the radius *R* will increase whereas the concentration *c* will decrease. This represents the relaxation of the deformed surface to a sphere (see SI). On the other hand, when the activity is large, i.e., *ζ* Δ*µ/β* ≫ *R*^−2^, the radius *R* decreases while the concentration *c* increases. The above equations imply that the surface constriction is governed only by the competition between activity dominates bending elasticity. However, note that we have ignored the effects of viscous dissipation and have implicitly assumed that the activity is high enough to have overcome these homogenizing effects.

In the regime where the activity dominates, i.e., *ζ* Δ*µ/β* ≫ *R*^−2^, we neglect the effects of bending elasticity and get

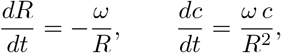

where *ω* = *ζ* Δ*µ/γ*. Solving for *R*(*t*) gives

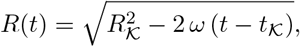

where *R*_𝒦_ is the radius of the cytokinetic ring at *t* = *t*_𝒦_. We emphasize that this is *not* the radius of the initial spherical surface, i.e., *R*_𝒦_ ≠ *R*_0_. For short times around *t*_𝒦_, a Taylor expansion leads to

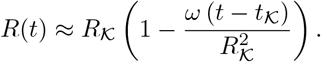

Using 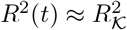 in the concentration equation, we get

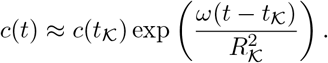

These solutions allows us to identify the constant ingression rate *u*_⊥_ and the exponential rate *k*_*m*_ of myosin accumulation:

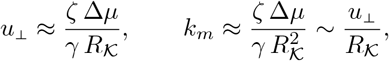

and predict a linear relationship between *k*_*m*_ and *u*_⊥_.

### Numerical Methods

To analyse the full nonlinear dynamics of our active geometrodynamic theory defined by the Equations (1)-(4), we use the finite-element method. We triangulate the surface and use the FEniCS library (Alnæs *et al*., 2015) to solve the resulting discrete system of equations. Time-marching was done using an implicit-explicit scheme.

## Notes

### Competing Interest Statement

The authors have declared no competing interest.

